# Mechanisms of 10-Hydroxyoctadecanoic acid resistance in *Streptococcus pneumoniae*

**DOI:** 10.1101/2025.06.06.658354

**Authors:** Cydney N. Johnson, Matthew W. Frank, Brendan T. Morrow, Qidong Jia, Christopher D. Radka, Jason W. Rosch

**Affiliations:** Department of Host-Microbe Interactions, St. Jude Children’s Research Hospital, Memphis, TN, USA; Graduate School of Biomedical Sciences, St. Jude Children’s Research Hospital, Memphis, TN, USA; Department of Microbiology, Immunology, and Molecular Genetics, University of Kentucky, Lexington, KY, USA

**Keywords:** hydroxy fatty acid, *Streptococcus pneumoniae*, glycosyltransferase, recombinase, membrane composition, cell charge, fatty acid resistance, phase variation, lipid profile

## Abstract

Profiles of human nasal colonization consistently demonstrate that *Staphylococcus aureus* and *Streptococcus pneumoniae* can co-exist in the nasopharynx. Several studies have demonstrated the antagonist relationship between the two organisms via several molecular mechanisms including competition for nutrients as well as via direct killing by hydrogen peroxide. During nasal colonization, the pneumococcus is in direct contact with the fatty acid *h*18:0, which is released into the extracellular environment by *S. aureus*. We report that *h*18:0 is specifically toxic to the pneumococcus amongst the pathogenic streptococci, providing a unique mechanism for interspecies competition during colonization. Exposure of cells to *h*18:0 revealed that *S. pneumoniae* could rapidly adapt to and overcome the observed toxicity. Whole genome analysis revealed the mechanism underlying this resistance being linked to a truncation of a glycosyltransferase in the capsule biosynthesis locus and a genomic inversion in the phase variation locus, leading to altered cell surface charge and membrane lipid composition. These physiological differences in the resistant isolates may aid in repelling toxic, charged fatty acids such as *h*18:0 from the cell membrane.

**IMPORTANCE:** The pneumococcus and *S. aureus* are two of the most well-characterized residents of the human nasopharynx; yet much remains unknown regarding how the two bacteria interact. Here, we describe the potential of *S. aureus–*produced *h*18:0, whose function and biological impact are still being described, to act as an inter-species competition molecule against *S. pneumoniae*, and how the pneumococcus can adapt to overcome its toxicity.

## INTRODUCTION

Bacterial oleate hydratase (OhyA) activity was first described in 1962, with its metabolic product identified as 10(*R*)-hydroxyoctadecanoic acid (*h*18:0) two years later^1,2^. OhyA genes are found in bacteria, but not mammals, and the encoded hydratases act on mammalian unsaturated fatty acids that contain either 9*Z* or 12*Z* double bonds^3–6^. These proteins are critical in detoxifying mammalian unsaturated fatty acids and promoting virulence^5,7,8^; specifically, the hydroxy fatty acids resulting from the OhyA reactions are not used by the bacteria but are rather released into the environment^7,8^. Recently, it has been shown that the major OhyA metabolite, *h*18:0, from *Staphylococcus aureus* stimulates a transcriptional cascade in macrophages that is driven by the activation of PPARα, leading to suppression of the innate immune response and increased expression of fatty acid oxidation genes to degrade the hydroxy fatty acid signals^9,10^.

*Streptococcus pneumoniae* (the pneumococcus) is a Gram-positive coccus that usually exists as a commensal of the human nasopharyngeal mucosa but is often characterized as a pathogen due to its ability to disseminate from the nasopharynx into the middle ear space, lower respiratory tract, blood, and brain^11^. Community-acquired pneumonia is the most common manifestation of invasive pneumococcal disease in those younger than five years or older than sixty-five years, despite the availability of antibiotics and vaccines^12,13^. All strains of the pneumococcus undergo phase variation, which is defined as the spontaneous, reversible phenotypic variation in colony opacity, from translucent to opaque^14^. Opaque variants have increased capsule production and decreased teichoic acid in the cell wall^15^. In mice, infection with opaque variants leads to increased morbidity, suggesting that there is a strong selection for organisms with the opaque phenotype during invasive infection^15^. Colonization is often associated with the translucent phenotype^16^. Phase variation in the pneumococci depends on DNA inversion events among three *hsdS* genes of a conserved type I RM system, known as the *cod* locus^17^, *ivr* locus^18^, or *spnIII*^19^. Genetic loci similar to *spnIII* have been identified in a diverse profile of bacterial genera^20–22^.

It has been known for decades that mammalian fatty acids, such as oleic acid (18:1), are toxic to streptococci, via destabilization of the bacterial membrane ^23,24^. To combat toxic fatty acids, bacteria encode a myriad of methods to deal with these pressures, including detoxifying the fatty acids via modification or directly incorporating them into their membranes^25^. *S. pneumoniae* encodes the fatty acid kinase system (FakAB) that allow for highly selective acquisition of host fatty acids found in human serum to replace *de novo* biosynthesis to promote membrane synthesis via scavenging from the host^26^. *S. pneumoniae* encodes three distinct FakB proteins (FakB1, FakB2, FakB3) that allow for the acquisition of saturated, monounsaturated, and polyunsaturated fatty acids, respectively^26^. More recently, it has been reported that the milk protein alpha-lactalbumin and its equine milk protein homologue, when in complex with 18:1, have bactericidal activity against pneumococci^27,28^. However, unlike many other Gram-positive bacterial pathogens as well as closely related *Streptococcus* species^5,29^, the pneumococcus does not encode an *ohyA* homologue to prevent the accumulation of the unsaturated, antimicrobial fatty acid 18:1.

The human nasopharynx is a diverse and complex niche. Upon approval and widespread use of the pneumococcal conjugate vaccine (PCV), the nasopharyngeal microbiome has undergone disruptions in the population structure of *S. pneumoniae*^30,31^. The change in colonizing pneumococci has likely altered this niche, also affecting *S. aureus*, another nasopharyngeal resident microbe^32–34^. Models of cocolonization with both species show that both species can co-exist in the nasal passages^35^, with stable dual-species biofilms able to be formed in the murine nasal passage^36^. Epidemiological studies report a recent increase in cocolonization rate of the human nasopharynx, with both species detected in upwards of 10% of patients^37,38^. However, the relationship between the two organisms seems antagonistic, and the molecular mechanisms by which they interact and compete for resources in this niche remain an active area of investigation. Studies have documented how the pneumococcus produces hydrogen peroxide to kill *S. aureus*^*39*^ and *S. aureus* produces catalase, which can be used to combat bacterial and host generated oxygen radicals^40^. In the female genital tract, OhyA produced by some lactobacilli sequesters 18:1 for phospholipid biosynthesis as *h*18:0 in an OhyA-dependent mechanism, enhancing bacterial fitness because only organisms that encode OhyA can utilize *h*18:0, while *Lactobacillus* species lacking OhyA are competitively disadvantaged ^41^. As a resident of the nasal microbiota, the pneumococcus is in direct contact and competition with bacteria such as *S. aureus* that encode an OhyA and release *h*18:0 into the environment^7,42^. We postulated that *h*18:0 released by *S. aureus* could be a potential mechanism by which this major human pathogen could eradicate competing bacterial species, specifically those lacking a homolog of OhyA such as *S. pneumoniae*.

Here, we report that *S. aureus*–derived *h*18:0 is toxic to the *S. pneumoniae*, a feature not widely shared by related pathogenic streptococci. Despite strong inhibitory activity, under selective pressure *S. pneumoniae* rapidly developed resistance to *h*18:0. Genetic analysis of resistant mutants uncovered the genetic basis for *h*18:0 fatty acid resistance which involved the mutation of a glycosyltransferase and a genomic rearrangement in the phase variation locus *spnIII* that is likely driven by a recombinase. The genetic changes in the resistant mutants altered both the surface charge of the resistant strains as well as modulated the lipid composition of their cellular membranes. These data underscore the potential role of *h*18:0 in mediating microbial competition during colonization, and bacterial strategies deployed to circumvent toxicity of this molecule.

## RESULTS

### 10-Hydroxyoctadecanoic acid is toxic to the pneumococcus

Oleic acid, the fatty acid 18:1, causes toxicity to several streptococcal species via membrane disuption^23,24^, though the extent to which this extends to the pneumococcus remains unknown.

*S. aureus* converts 18:1 to *h*18:0 via OhyA; however, *S. pneumoniae* lacks this protein. We sought to determine whether the host-derived precursor and the bacterially modified fatty acids produced from the *S. aureus* OhyA reaction are toxic to pneumococci. *S. pneumoniae* is composed of over 100 different serotypes that are determined by the polysaccharide capsule produced by the bacteria. Pneumococcal strains TIGR4 (serotype 4), BHN97 (serotype 19F), D39 (serotype 2), Serotype 7F, A66.1x (serotype 3), and Tupelo (serotype 14) were grown in C+Y media supplemented with solute alone (DMSO) and either 10uM 18:1 or 10µM *h*18:0. There was considerable heterogeneity in how different strain backgrounds responded to the supplemented fatty acid (Fig. 1). All strains tested were susceptible to 18:1 toxicity except

**Figure 1.**
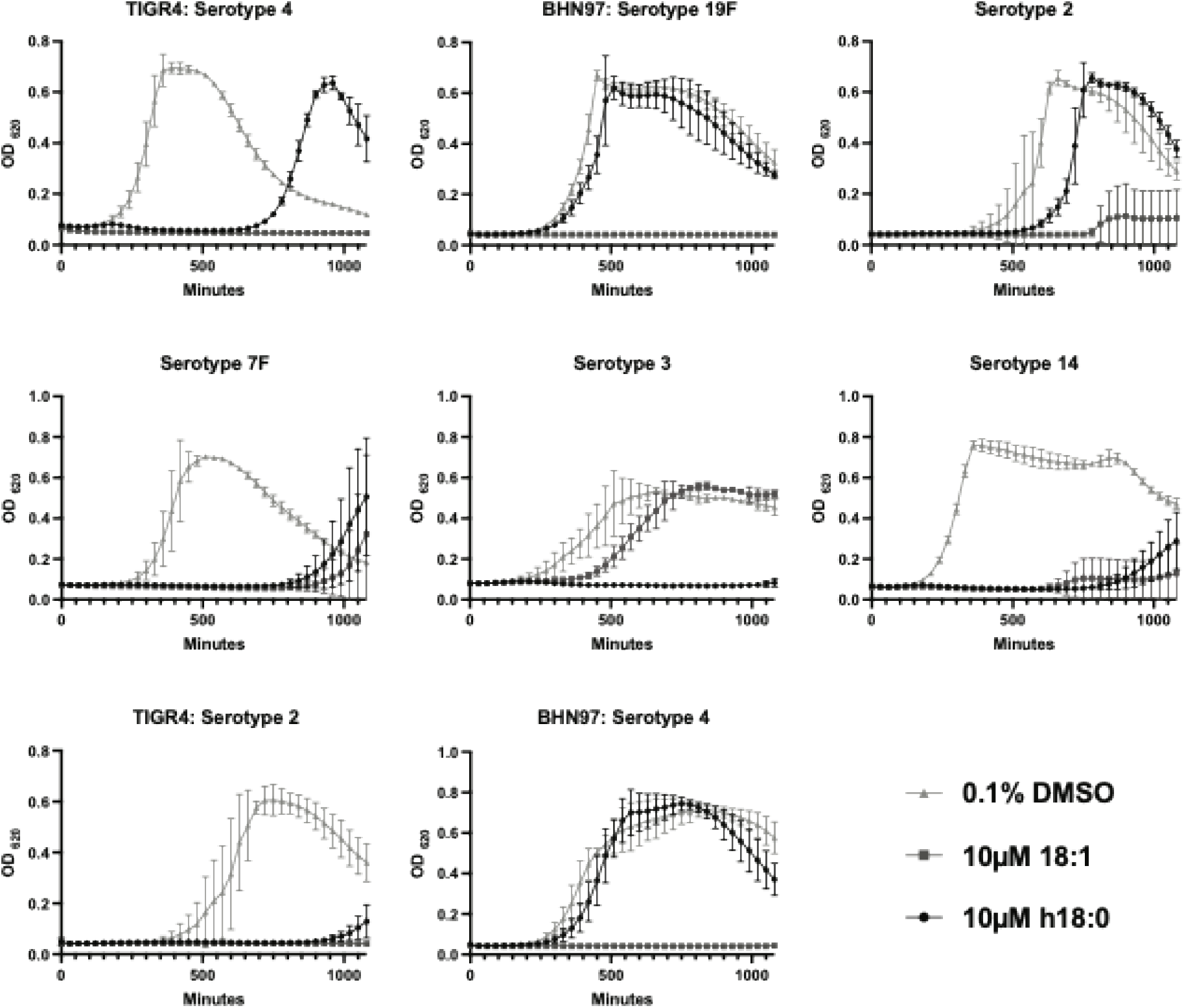
*h*18:0 exhibits bactericidal activity against pneumococcal capsule types that are associated with invasive disease. Pneumococci were grown in complex media with 0.1% DMSO, 0.1% DMSO + 10 µM 18:1, or 0.1% DMSO + 10 µM *h*18:0. Growth was evaluated by measuring optical density every 30 minutes for 20 hours. Serotypes are denoted above each graph. All experiments were done in biological triplicate with technical triplicates, except 7F and A661.x, which were performed in biological duplicate with technical triplicates. Mean and standard deviation values are shown.

A661.x (serotype 3), which was inherently resistant, confirming the broadly inhibitory nature of 18:1 which has been seen in several bacterial species^23–25,41^.

The susceptibility of strains to *h*18:0 susceptibility was more heterogeneous across strains and capsule backgrounds. D39, like TIGR4, was susceptible to *h*18:0 toxicity but established normal growth after a prolonged lag phase. Strains 7F, A66.1x, and Tupelo largely remained susceptible to *h*18:0 toxicity; however, these strain backgrounds typically did not exit lag phase until much later in the assay.

The pneumococcal capsule is a highly diverse structure in terms of both composition as well as charge, both of which play important roles in bacterial fitness under various conditions. To determine whether the phenotypes rely upon the differences between capsule types, we generated capsule switched variants of TIGR4 expressing the 19F capsule and BHN97 expressing the type 4 capsule. The capsule-switched strains maintained the susceptibility profile of the strain’s parental expressing their native capsule (Fig. 1), indicating that differing susceptibility to 18:1 and *h*18:0 was not related to the different polysaccharide capsules being produced but rather another genetic determinant.

### 10-Hydroxyoctadecanoic acid toxicity is specific to closely related *Streptococcus* species

*S. pneumoniae* does not encode a gene producing OhyA but other closely related Gram-positive bacteria do. According to published genomes, *S. pyogenes* (KEGG T00050, Spy_0470) and *Enterococcus faecalis* (KEGG T00123, EF3303), once classified as group D streptococci; *Streptococcus mutans* (KEGG T00100, SMU_515 and SMU_1584c); and *Streptococcus agalactiae* (KEGG T00091, SAG1508) all encode for a predicted functional oleate hydratase. *S. pyogenes* is known to be susceptible to 18:1 toxicity^24^, but the susceptibility of the other species has not been determined, so we next asked whether these closely related bacteria are susceptible to 18:1 and *h*18:0 toxicity (Fig. 2). As expected, *S. aureus* was resistant to 18:1 and *h*18:0 toxicity in an OhyA-independent manner (Fig. 2A, B). *E. faecalis, S. mutans*, and *S. agalactiae* were not affected by either 18:1 or *h*18:0 (Fig, 2C, D, E). *S. pyogenes* was susceptible only to *h*18:0 (Fig. 2F). These data underscore that even amongst closely related streptococcal species, sensitivity of potential antimicrobial fatty acids varies considerably.

**Figure 2.**
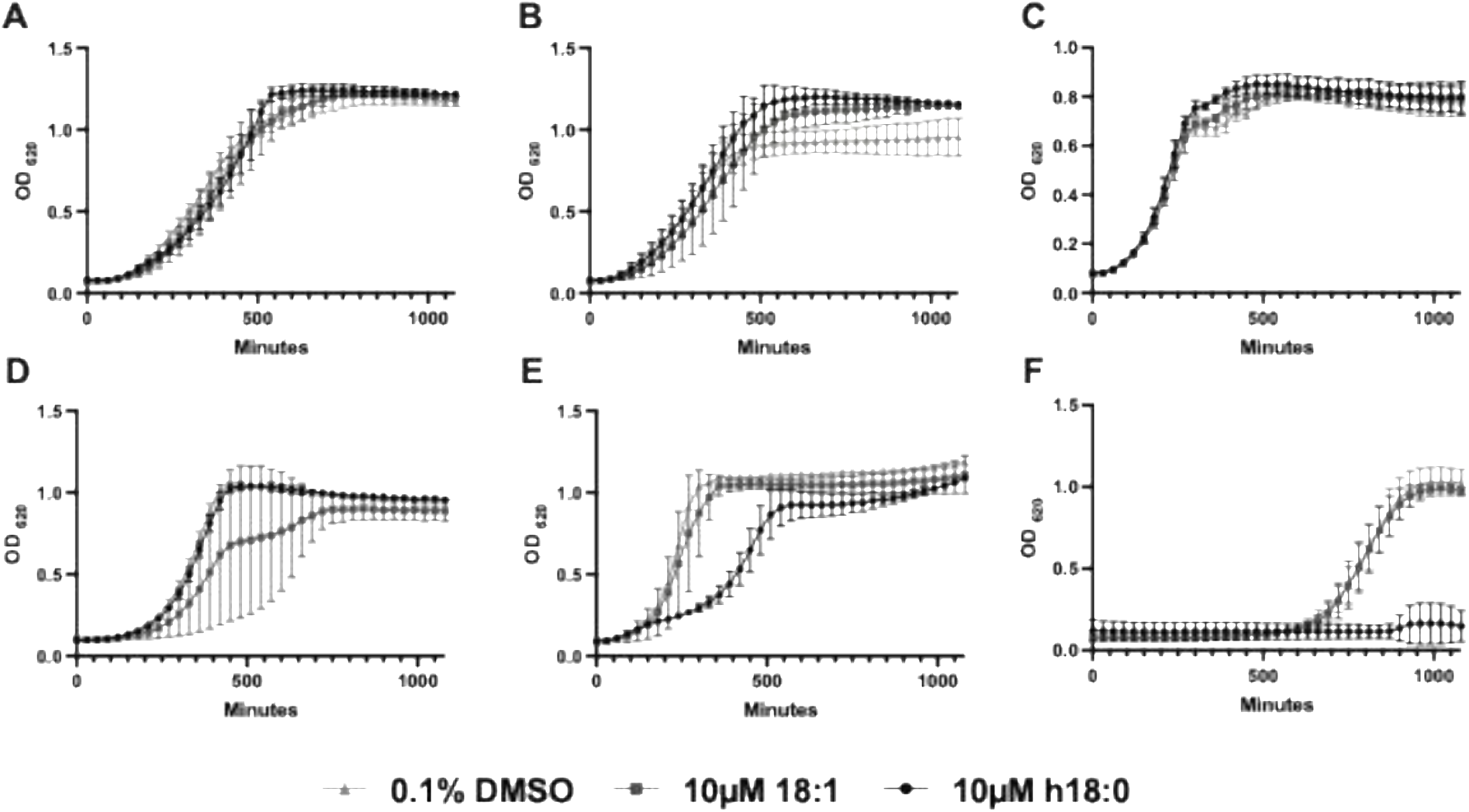
*h*18:0 toxicity is not conserved across Gram-positive species. Bacterial strains were grown in media supplemented with 0.1% DMSO, 0.1% DMSO + 10 µM 18:1, or 0.1% DMSO + 10 µM *h*18:0. All experiments were done in biological triplicate with technical triplicates. (A) *S. aureus*, (B) *S. aureus* Δ*ohyA*, (C) *E. faecalis*, (D) *S. mutans*, (E) *S. agalactiae*, (F) *S. pyogenes*. Mean and standard deviation values are shown.

### Long-read sequencing reveals that resistance to 10-hydroxyoctadecanoic acid relies on a recombinase and glycosyltransferase

As noted in the initial growth kinetics, following a prolonged phase most strains of *S. pneumoniae* began to replicate at normal growth rates. These data suggest that during this prolonged exposure that resistant mutants likely arose in the population that were resistant to the inhibitory activity of *h*18:0. These clones demonstrated normal growth kinetics in standard C+Y media that was the base media for all experiments (data not shown). Supplementation of media with *h*18:0 confirmed that all three clones demonstrated increased resistance to the inhibitory activity of *h*18:0 (Fig. 3A) indicating a heritable resistant mechanism.

**Figure 3.**
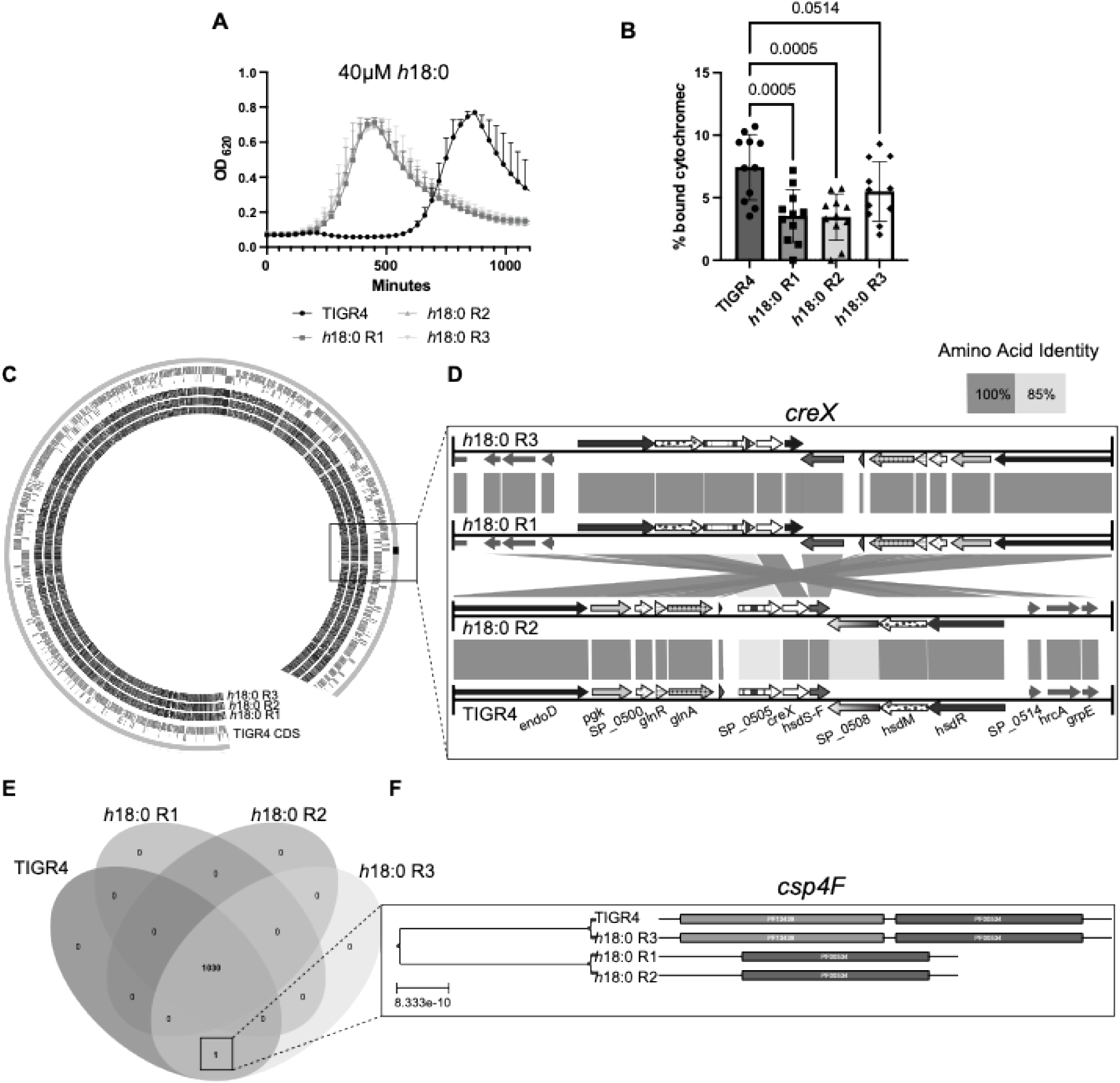
Mutation of a glycosyltransferase or genetic recombination can cause *h*18:0 resistance. (A) *h*18:0-resistant mutants can grow in 40 µM *h*18:0; wild-type TIGR4 is susceptible to the toxicity. Mean and standard deviation values are shown. (B) Resistant mutants exhibit a more positive cell charge due to decreased binding of cytochrome *c*, a positively charged protein. Mean and standard deviation values are shown. Statistical significance was determined by one-way ordinary ANOVA. (C) Short-and long-read sequencing was used to *de novo* assemble genomes for wild-type TIGR4 and *h*18:0-resistant mutants (inner rings); genomes were analyzed by using zDB^46^. Individual CDS represented in the TIGR4 genome. The outermost ring represents contigs from the reference assembly. Inner rings represent sequence presence of protein homologs compared to wild-type TIRG4. (D) Genomic analysis of the locus around *creX* reveals an inversion (crossed amino acid identity lines between genomes) and mutation of genes flanking *creX*. Homologous genes display identical patterns. Lines between genes represent amino acid similarity. (E) Venn diagram of KO groups among the genomes. (F) *cspF4* domain visualization among the four genomes. * p ≤0.05, ** p ≤0.01, *** p ≤0.001, **** p ≤0.0001.

We next sought to determine if the resistant clones demonstrated physiological characteristics that could explain the observed resistance patterns against *h*18:0. Hydroxy fatty acids are typically negatively charged due to the carboxylic acid group. While our capsule switched mutants did not demonstrate appreciable difference in *h*18:0 sensitivity, we postulated that capsule-independent modulation of surface charge could be involved in resistance. This was measured using a cytochrome *c* binding assay, which is widely utilized to measure relative surface charge^43–45^. We found that these clones exhibited a more positively charged cell surface, as determined by cytochrome *c* binding, compared with the sensitive parental strain (Fig. 3B).

To determine the genetic basis of this resistance, the *h*18:0-resistant mutants (strains R1, R2, R3) were subjected to both short-read Illumina and long-read PacBio sequencing. Genomes were assembled *de novo* and analyzed using zDB^46^ (Fig. 3C). We observed a mutational hotspot surrounding the site-specific recombinase–encoding gene *creX* (SP_0506) in the *h*18:0-resistant mutants (Fig. 3D). The gene *creX*, along with the restriction modification system *hsdM* and *hsdR*, is part of the *spnIII* locus, which is highly conserved across pneumococcal strains and is involved in phase variation^19^. Specifically, *h*18:0 R1 and R3 have undergone inversions of this locus that centers around *creX*. Unlike the other resistant isolates, *h*18:0 R2 had mutations in the genes SP_0505 and SP_0508, which are up-and downstream of *creX*, respectively. These genes encode proteins that are about 85% identical at an amino acid level to the TIGR4 proteins. SP_0505 and SP_0508 (*hsdS*) are putative subunit S proteins of type I restriction-modification systems. CreX driven recombination of *hsdS* is a known rearrangement that leads to phase variation in pneumococci^47^.

We also queried for additional genetic features unique to the *h*18:0-resistant clones. Specifically, we utilized the KEGG database to identify any pathways present or absent in our *h*18:0-resistant clones, then created a Venn diagram illustrating the distribution of unique orthologs between genomes (Fig. 3E). There was only a single hit in *csp4F* (SP_0351), a glycosyltransferase in the capsule biosynthesis locus (Fig. 3F), with *h*18:0-resistant clones R1 and R2 deleting the 5′ end of this gene. Despite these mutations, these strains still produced type 4 capsule as determined by latex agglutination and ELISA (data not shown). These findings suggest that changes in the phase variation locus *spnIII* and in capsule glycosylation may promote *h*18:0 resistance in *S. pneumoniae* TIGR4.

### *S. pneumoniae that are* resistant to 10-hydroxyoctadecanoic acid have altered membrane composition

Previous studies have implicated changes in lipid composition when bacteria are grown with fatty acids^48–50^. Phase variation affects lipid and cell wall composition^51,52^; therefore, we determined whether our *h*18:0-resistant clones with genomic rearrangements in this *spnIII* locus involved in phase variation have altered lipid profiles in their cell membranes. Utilizing gas chromatography, we quantified the fatty acid composition of the lipid membrane in wild type TIGR and the h18:0-resistent clones (Fig. 4A). The *h*18:0-resistant clones had significantly less 18:1 and more 18:0 in their lipids compared to WT. The h18:0-resistant clones contained significantly more saturated and fewer unsaturated fatty acids in their membrane than WT clones did (Fig. 4B). Additionally, the phosphatidylglycerol (PG) profile for wild type and resistant strains were analyzed by mass spectrometry. The different PG species can contain in total 0, 1, or 2 double bonds in their fatty acid combinations. In the total PG, the h18:0-resistant strains contained a lower percentage of PG species that contained 2 double bonds like 36:2 and an increased percentage with 0 or 1 double bonds (Fig 4C) Using [1-^14^C}acetate labeling, we characterized the lipid profile of TIGR4, the *h*18:0-resistant clones, and BHN97 (Fig. 4D). The resistant clones have significantly increased amounts of diacylglycerol and glucosyl-diacylglycerol, while phosphatidylglycerol and galactosyl-glucosyl-diacylglycerol amounts are decreased which is similar to BHN97. In conclusion, *h*18:0-resistant clones have different lipid compositions in both the fatty acid composition of their lipids and the amount of the different lipid species than wild-type *S. pneumoniae*.

**Figure 4.**
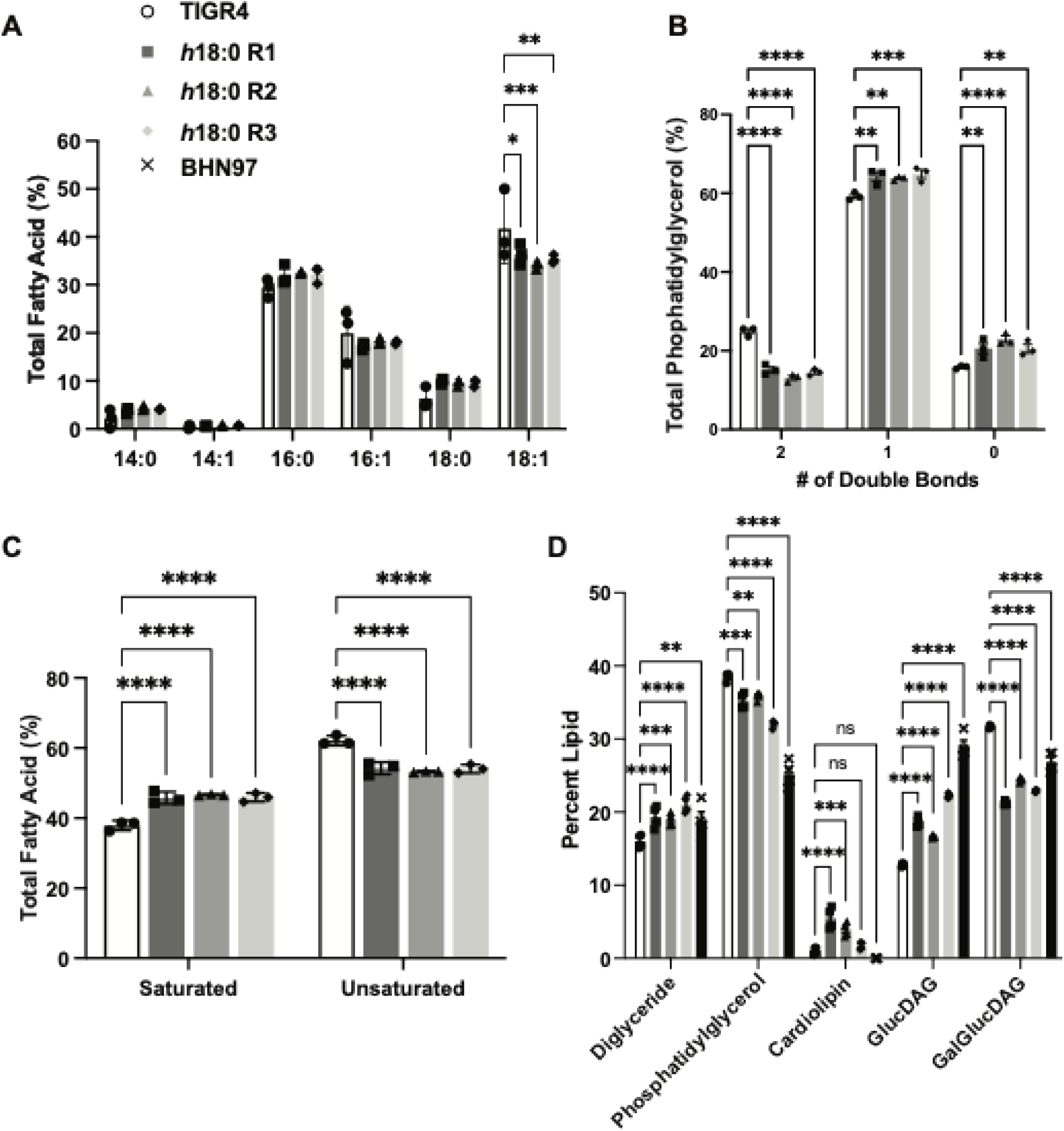
Pneumococci resistant to *h*18:0 have altered membrane lipid composition. (A) Fatty acid composition of the membrane lipids composition was determined for wild-type and *h*18:0-resistant TIGR4 strains. (B) PG species are grouped by the number of double bonds they contain in the fatty acids. (C) From the FAME analysis, fatty acids are grouped together as saturated or unsaturated fatty acids. (D) Acetate labeling was performed to determine the lipid composition of the cell membranes. Mean and standard deviation values from biological triplicates are shown for all panels. Statistical significance was determined by one-way ordinary ANOVA. * p ≤0.05, ** p ≤0.01, *** p ≤0.001, **** p ≤0.0001.

## DISCUSSION

The human nasopharynx is a diverse and complex niche that promotes constant interactions between bacteria to compete for limited space and nutrients. The pneumococcus and *S. aureus* are two of the most well-known residents of the human nasopharynx; yet much remains unknown regarding how the two bacteria interact. *S. aureus* oleate hydratase OhyA converts host 18:1 to *h*18:0^8^. *h*18:0 is released from the bacterial cells as a free fatty acid and would, therefore, be presented to the pneumococcus as an unesterified fatty acid. As of yet, no evidence indicates that host cells can integrate *h*18:0 into a lipid, raising the question as to what the impact of this fatty acid is on the neighboring microbiota. In this study, we discovered that *h*18:0 is toxic to the pneumococcus, supporting the idea that *h*18:0 may serve as a secreted metabolite used by *S. aureus* to mediate competition amongst other bacterial species.

While we observed that *h*18:0 effectively inhibited growth of most pneumococcal strains, we noted that resistance consistently and readily emerged against this antimicrobial fatty acid. While most pneumococcal genetic backgrounds and serotypes were sensitive to *h*18:0, there were notable exceptions. For example, serotype 3 was the only tested pneumococcal serotype with complete resistance to *h*18:0. Serotype 3 infections are characterized as having severe clinical manifestations and causing invasive disease, including empyema, bacteremia, cardiotoxicity, and meningitis, with a fatality rate over 30%^53^. The *h*18:0 resistance by serotype 3 may be due to the highly mucoid capsule that this serotype produces. After generating TIGR4 clones resistant to *h*18:0, subsequent analysis of the lipid profile of the *h*18:0-resistant mutants revealed subtle but consistent alterations in lipid membrane composition. The *h*18:0-resistant clones contained significantly more saturated and fewer unsaturated fatty acids in their membrane than the parental wild type strains. In addition, the resistant clones have significantly increased amounts of diacylglycerol and glucosyl-diacylglycerol, while phosphatidylglycerol and galactosyl-glucosyl-diacylglycerol amounts are decreased, like BHN97. This is in agreement with the measurement of surface charge differences in the resistant mutants, suggesting electrostatic repulsion may confer a protective benefit against the toxicity of *h*18:0. There is also the noteworthy observation that the *S. pneumoniae* does not incorporate *h*18:0 into their membranes, meaning either their FakB proteins do not accept *h*18:0, or their lipid biosynthetic machinery does not accept *h*18:0 acyl chains. Future studies are necessary to clarify the mechanistic relationship between membrane lipid composition and fatty acid resistance.

The genetic basis of fatty acid resistance has been difficult to determine over the last few decades. Many of the studies have utilized sequencing to examine a selected subset of genes

—generally, the Fak locus because inactivation of genes in this pathway can lead to fatty acid resistance^54^— and have found single mutations that lead to fatty acid resistance^55^. In this study, we used whole-genome sequencing for our resistant clones to take an unbiased approach to discovering genetic mechanisms of *h*18:0 resistance. By using both short- and long-read sequencing, we were able to generate *de novo* genomes of our *h*18:0-resistant clones with sufficient coverage and depth. This led to the elucidation that in *S. pneumoniae* TIGR4, *h*18:0 resistance can come from multiple mechanisms. Two of the three clones had large genomic inversions surrounding *creX. creX* is part of the phase variation–determining locus *spnIII*, which is highly conserved across pneumococcal strains; genomic recombination of this locus alters locus expression and controls phase variation^19^. The zDB analysis only enabled analysis of the 20 kB surrounding this gene, so the inversion may have been larger than what we observed. Intriguingly, it has been reported that some serotype 19F strains lack *creX*^56^, which may explain why BHN97 is resistant to *h*18:0 independent of the capsule type that is expressed.

Truncation of *cps4F* in *h*18:0-resistant clones R1 and R2 would presumably lead to an inactive glycosyltransferase. In a study surveying pneumococcal isolates from Malaysia, *cps4F* was conserved across the sequenced serotype 4 isolates^57^. Despite these mutations, the strains expressed serotype 4 capsule as determined by latex agglutination assays. Differences in glycosylation patterns have the potential to change cell surface potential, which may aid in repelling charged fatty acids from the cell membrane. Increasing cell charge inhibits the ability of fatty acids to permeate the cell wall, which will inhibit access to the sites of action on the inner membrane^42,58,59^. Despite having limited genetic mutations, all the *h*18:0-resistant mutants had altered composition of lipid species in their cell membranes. Future studies exploring the mechanism of resistance in BHN97 may uncover additional mechanisms of fatty acid resistance.

## MATERIALS AND METHODS

### Bacterial strains and growth conditions

Pneumococcal strains were grown on tryptic soy agar (Sigma Aldrich) supplemented with 3% defibrinated sheep blood and 20 μg/mL neomycin.

*S. aureus* AH1263^60^ and AH1263 Δ*ohyA*^8^, *E. faecalis* OG1RF, *Streptococcus mutans* UA159, *Streptococcus agalactiae* 2603, and *Streptococcus pyogenes* HSC5 were grown in Todd-Hewitt media (BD) supplemented with 2% w/v yeast extract (Gibco). Pneumococcal cultures were inoculated from newly streaked TSA blood agar plates into C+Y, a semi-synthetic casein liquid media with 0.5% w/v yeast extract^61^, and grown at 37°C, 5% CO_2_. Growth curve measurements were read in a 96-well plate in a Biotek Synergy, with starting OD_620_ of 0.05-0.1 for all strains.

### Cytochrome *c* binding assay

The cytochrome *c* binding assay was modeled after previously published assays^45,62^. Pneumococci were grown to an OD_620_ of 0.4 in 7 mL C+Y. 1 mL of the culture was pelleted at 13,000 rpm for 1 minute and resuspended in 400 µL of 1 M sterile 4-(2-hydroxyethyl)-1-piperazineethanesulfonic acid (HEPES, Sigma Aldrich). Cytochrome *c* (Sigma) was added to a final concentration of 0.5 mg/mL; the cell mixture was incubated at room temperature for 10 minutes, followed by centrifuging the bacteria at 13,000 rpm for 1 minute.

The assay was performed in biological triplicate, and the experiment was repeated three times. Cytochrome *c* remaining in the supernatant was quantified by measuring absorbance at 535 nm wavelength and compared to that of samples with no bacteria.

### DNA extraction, sequencing, and genomic assembly

DNA was extracted via phenol/chloroform extraction from wild-type *S. pneumoniae* TIGR4, and the three independently derived *h*18:0-resistant mutants were grown to stationary phase in either 0.01% DMSO (wild-type) or 0.01% DMSO + 10 µM *h*18:0 (resistant mutants). High molecular weight DNA was submitted for Illumina short-read sequencing on the HiSeq platform and PacBio Revio long-read sequencing performed by the Hartwell Center at St. Jude Children’s Research Hospital.

### Mass Spectrometry of Phosphatidylglycerol

Wild-type and *h*18:0-resistant TIGR4 strains were grown in C+Y media for 6 hours; after which, cells were pelleted, and the supernatant was removed. Cell pellets were stored at −20°C until analysis. Lipids were extracted from the cells by using the Bligh & Dyer method^63^. Lipid extracts were resuspended in chloroform:methanol (1:1). PtdGro was analyzed by using a Shimadzu Prominence UFLC attached to a QTrap 4500 equipped with a Turbo V ion source (Sciex). Samples were injected onto an Acquity UPLC BEH HILIC, 1.7 µm, 2.1 × 150-mm column (Waters) at 45°C with a flow rate of 0.2 mL/min. Solvent A was acetonitrile, and solvent B was 15 mM ammonium formate, pH 3. The HPLC program was the following: starting solvent mixture of 96% A/4% B, 0 to 2 min isocratic with 4% B; 2 to 20 min linear gradient to 80% B; 20 to 23 min isocratic with 80% B; 23 to 25 min linear gradient to 4% B; 25 to 30 min isocratic with 4% B. The QTrap 4500 was operated in the Q1 negative mode. The ion source parameters for Q1 were as follows: ion spray voltage, −4500V; curtain gas, 25 psi; temperature, 350°C; ion source gas 1, 40 psi; ion source gas 2, 60 psi; and declustering potential, −40V. The system was controlled, and data analyzed by the Analyst software (Sciex).

The sum of the areas under each peak in the mass spectra were calculated, and the percentage of each molecular species present was calculated with LipidView software (Sciex).

### Fatty Acid Analysis by gas chromatography

Fatty acid methyl esters were prepared from the lipid extracts using anhydrous methanol/acetyl chloride. The fatty acid methyl esters were analyzed by a Hewlett-Packard model 5890 gas chromatograph equipped with a flame ionization detector and separated on a 30 m × 0.536 mm × 0.50 µm DB-225 capillary column (Agilent). The injector was set at 250°C, and the detector, at 300°C. The temperature program was as follows: initial temperature of 70°C for 2 min, rate of 20°C/min for 5 min (final temperature, 170°C), rate of 2°C/min for 10 min (final temperature, 190°C), hold at 190°C for 5 min, rate of 2°C/min for 15 min (final temperature, 220°C), hold at 220°C for 5 min. The identity of each fatty acid methyl ester was determined by comparing its retention time with fatty acid methyl ester standards (Matreya). The composition was expressed as weight percentages.

### [1-^14^C] Acetate Incorporation into Lipids

When strains grown in C+Y media under standard growth conditions reached an OD_620_ of 0.2, 20 µCi of [1-^14^C] acetate was added and incubated at 37°C to an OD_620_ of 0.8. Cells were collected by centrifugation and washed twice with PBS and once with water. Lipids were extracted using the Bligh and Dyer method. Radiolabeled lipids were quantified by scintillation counting. The lipids were separated by loading equivalent amounts of radioactivity onto a silica gel H thin-layer plate and were developed in ethanol:chloroform:triethylamine:water:0.5M EDTA (34:30:35:6.5:0.210, vol/ vol/vol/vol/vol). The radiolabeled lipids were visualized using the Typhoon FLA 9500 (GE Healthcare). Lipids were identified with known standards.

## ACKNOWLEDGEMENTS

We would like to thank Cherise Guess, PhD, ELS of the Department of Scientific Editing at St. Jude Children’s Research Hospital for providing edits to the manuscript. This research included experiments conducted by the Hartwell Center for Bioinformatics & Biotechnology, which is supported, in part, by ALSAC and National Cancer Institute grant P30 CA021765. C.N.J. is supported by NIH grants T32-AI106700 and F32-AI286447. C.D.R. is supported by NIH grant R00-AI166116. The content of this article is solely the responsibility of the authors and does not necessarily represent the official views of the National Institutes of Health.

